# How perception and attention participate in reading skill?

**DOI:** 10.1101/2021.06.07.447429

**Authors:** Sunita Gudwani, Manju Mehta, Rajesh Sagar, Madhuri Behari, Vaishna Narang, Sadanad Dwivedi, N.R. Jagannathan, S. Senthil Kumaran

**Affiliations:** Escorts Heart Institute and Research Centre, India; Child Psychology Unit, Department of Psychiatry, All India Institute of Medical Sciences, New Delhi, India; Department of Psychiatry, All India Institute of Medical Sciences, New Delhi, India; Department of Neurology, Fortis Hospital, Vasant Kunj, New Delhi, India; Department of Linguistics, School of Language Jawahar Lal University, New Delhi, India; Department of Biostatistics, All India Institute of Medical Sciences, New Delhi, India; Department of NMR & MRI Facility, All India Institute of Medical Sciences, New Delhi, India

**Keywords:** Visuospatial attention, executive-control, dorsal-ventral stream, symbol-string, prelexical-lexical perception

## Abstract

Reading creates accessibility to the outer world information and is an integral skill for academic achievement. It involves visual perception, reasoning at symbolic level (text reading) and phonological processing. But does perception, visuospatial processing at pre-symbolic level affect reading skill? To understand and untangle the mechanism fifty children with reading problem (developmental dyslexia, DDC) and twentyfive age-matched typical readers (healthy controls, HC) were studied. In DDC, the variable-performance of non-symbolic visual search (pictorial-level) was associated with bilateral brain activity (functional MRI) of ventral stream (inferior occipital and fusiform) and greater involvement of frontal-prefrontal regions, suggesting the modified dorsal route-gating for figure-ground filtering. Performance variability in picture-concept reasoning (71.9% and significant group mean difference) and visuospatial-organization (51.3%) of DDC children indicate attentional or executive differences at nonsymbolic level. In symbolic discrimination performance, no significant difference observed for single alphabet or number (digit) reading but significant differences at word level for visual and phonological performance (considering orthographic differences), attributes to distractor-inhibition problems. Nonsymbolic to symbolic coding/decoding speed and writing error (copying) differences are suggestive of execution variability rather than motor coordination in DDC. In HC, the top-down dorsal route gating and hypoactivity induced in right frontal-anterior cingulate cortices, play a major role in reading by allocating spatial attention and inhibiting distractors. It has been elaborated in the “neurobiological model”.

**How perception and attention participate in reading skill?:** *Research Highlights:*

- Visuospatial attention, executive control (nonsymbolic level) and visual discrimination (symbolic level) performance contribute to reading.
- In children with reading problems (dyslexia) pictorial visual processing task (fMRI) recruited higher activation in right hemispheric frontal, inferior parietal and anterior cingulate regions.
- Bilateral activation of occipital-striate, extra-striate, fusiform for nonsymbolic is associated with significantly more errors at bigger-symbol-string, in dyslexia.
- Auditory discrimination differences in shallow-deep orthography.
- Correlation of non-symbolic executive and symbolic performances.
- Role of the prelexical visual perception and dorsal-route gating in reading skill explained as neurobiological model.

## 1. Introduction

Reading, a cultural trait, is a skill utilizing cognitive, perceptual and motor aspects of the neural mechanisms that had evolved for other purposes (Benischek et al, 2020; Valdois, Lassus-Sangosse, and Lobier, 2012; Vidyasagar and Pammer, 2010). Efficient mapping of the horizontally arranged symbols (alphabetic system) (Ahissar, et al, 2015; Doron, et al, 2012), has been reported to begin primarily as a phonological process and as the strongest predictor of reading skill (Snowling et al, 2020; Ziegler, Pech-Georgel, George, Lorenzi, 2009; Ziegler and Goswami, 2005). Reading is reception, discrimination, identification and labeling of symbols. Unfolding perception mechanism in reading, it is essential to understand the continuum from non-symbolic to symbolic in typical (normal) and non-typical (dyslexic) readers.

Developmental dyslexia, a learning difficulty of reading was first described more than a century ago (Morgan, 1896). It is recognized as the most common of developmental disorders, two thirds of all specific learning difficulties (Shaywitz and Shaywitz, 2005) and has a prevalence of 5-17% in school going children (Fletcher et al, 2007; Lyon et al, 2003; WHO, 1992). According to ICD-10 (F81.0, F81.1, F81.2, and F81.3), it is defined as difficulty in reading and writing skill acquisition (World Health Organization 1992: p. 245) that are not accounted by mental age, intellectual ability, visual acuity or schooling (Pennington, 2009; American Psychiatric Association, 1994). Etiology of this complex reading problem is disputed underlying mechanism(s) (Catts, 2021; Zoccolotti, de Jong, and Spinelli, 2016) involving visual attention, visuo-spatial working memory, executive functions, grapheme-phoneme mapping (orthographic-phonological), auditory-verbal decoding, lexico-semantic, syntactic, and comprehension processing (Ahissar, 2017; Ahmad, 2014; Arrington et al, 2019; Ashkenazi et al, 2013; Catts, 2021; Jaffe-Dax, et al, 2016; Martini et al, 2015; Pammer, 2014; Sela et al, 2012; Shaywitz and Shaywitz, 2005; Siok et al, 2009; Snowling et al, 2020; Truong et al, 2014; Vidyasagar and Pammer, 2010). In dyslexia performance inconsistencies across different tasks are unable to explicitly indicate an underlying cause (Catts, 2021; Ramus and Ahisaar, 2012; Roach and Hogben, 2007), since a single impairment may underlie broad range of difficulties in the perception dynamics (Ahissar, 2007), hence dyslexia is conceptualized as an amodal sluggish attentional shift (Krause, 2015; Ramus and Ahisaar, 2012; Roach and Hogben, 2007; Stenneken et al, 2011; Zoccolotti et al, 2016).

In visual perception, retina continuously encodes the vast cluttered information of natural visual scenes that pose processing limitation. For optimal interaction with surroundings only a subset information or the relevant aspects of visual scene needs to be prioritized and reach the awareness (Kastner and Pinsk, 2004; Roach and Hogben, 2007). In brain, primary visual perceptions depend upon intact early visual areas and specific higher-order parietal areas (Pollen, 2011) with the visuospatial network (Banker et al, 2020; Diehl et al, 2014), visual filtering of posterior parietal cortex (Friedman-Hill et al, 2003). Magnocellular system plays a role in visual discrimination (Ramus and Ahissar, 2012; Stein, 2014) where fast visual scene changes are processed (Livingstone et al, 1991) by inhibiting the parvocellular system (Stein and Walsh, 1997), or magnocellular being inhibited by the parvocellular system (Burr et al, 1994). However, in either case, the dyslexic deficits could not be explained (Zoccolotti et al, 2016). In dyslexia, processing speed, attentional focus modulation (Roach and Hogben, 2007; Vidyasagar and Pammer, 2010), decision making based on available visual information (Eckstein et al, 2004) and non-verbal cognitive loops (Margolis et al, 2018) may be impaired. Both, the dorsal and ventral pathways (Gallivan, et al, 2011; Zhou et al, 2015) or only the dorsal pathway could be affected (Stein and Walsh, 1997; Vidyasagar and Pammer, 2010). In reading, selective ‘visual attention’ and filtering of irrelevant information plays a critical role (Ding et al, 2016; Roach and Hogben, 2007). When considering symbolic system of reading, dyslexia has been associated with visual pattern recognition, auditory perception and phonological discrimination deficits due to dysfunctional occipito-temporo-frontal connections (Boets, 2014; Brem et al, 2020; Ding et al, 2016; Goswami et al, 2011; Lipowska et al, 2011; van der Mark et al, 2011), it is unclear whether they co-occur or actually characterize different subtypes of dyslexia (Ramus and Ahissar, 2012). The orthographic depth of language also influences cortical activity that further complicates the measurement of dyslexic differences (Martin, Kronbichler, and Richlan, 2016; Richlan, 2014).

To explore the role of pictorial, visuospatial and auditory processing in reading, the present study investigated non-symbolic to symbolic perceptual reasoning including experiment-1 with two parts (a) for visual perception (picture completion); and (b) for picture-concept the visuospatial reasoning (object assembly) and executive control (block design sub-test); experiment-2 for nonsymbolic to symbolic interface with (i) visual pattern recognition (ranging from non-symbolic shapes to symbolic), (ii) attention and judgment (coding icon-digit symbol); (iii) reading (alphabets, numbers, words as meaningful strings) in both deep and shallow orthographies (English and Hindi respectively); experiment-3 symbolic visuomotor execution (copying, writing-alphabets, numbers, words) and experiment-4 for phonological awareness (auditory discrimination of minimal pairs in two languages). These aspects of perceptual execution were investigated in typical (healthy normal) and non-typical (dyslexic) readers by performance assessment and cortical activity (functional MRI).

## 2. Methodology

The study was designed as a cross-sectional observation. After the Institute Ethical Sub-committee approval, total of 132 subjects were screened and fifty three (53) were recruited at out-patient clinic (specific learning disorder-SLD) of Psychiatry department and neuroimaging was done at department of Nuclear Magnetic Resonance Imaging. All the parents’ signed informed consent for their child participation. Inclusion criteria for children allocated to impaired reading group were 8 to 15 years of age, right handed (Edinburgh Inventory, Oldfield, 1971), IQ >80, normal/ corrected vision, bilingual, biliterate (Hindi and English) and diagnosed with developmental dyslexia. The subjects were clinically diagnosed based on ICD-10 and DSM-5 signs of reading rate, reading errors, reading comprehension, writing skill, writing rate, letter shape, spelling errors, phonological errors, mathematical calculations, etc. Exclusion criteria were comorbidity of psychiatric disorder, neurologic lesion, attention deficit hyperactive disorder and contraindication for magnetic resonance imaging (MRI). Inclusion criteria for age-matched, healthy controls were age 8 to 15 years, right handed, IQ >80, normal/ corrected vision, bilingual, biliterate (Hindi and English) and average errors (within one Standard Deviation) in reading, writing, spelling, good academic records. Exclusion criteria for healthy controls were similar to dyslexic children. Three dyslexic children and one healthy subject were excluded due to incidental neurological findings in MRI (lesion).

The clinical assessments were done in a non-distracting comfortable room. Semi-structured interview with parents and subject included demographic details with (a) socio-economic status, financial expenditure on child’s education; (b) parental education, schooling, medium of education, occupation (c) family history of motor, speech and/or language disorders/delay, any known genetic disorder, mental illness; (d) history of child’s development (including motor coordination), general health, interpersonal relationships, language acquisition, proficiency, preference, etc. Clinical assessments consisted of Malin’s Intelligence Scale for Children (MISIC, Malin, 1969; Regional adaptation of Wechsler Intelligence Scale for children), and Battery for diagnosis of specific learning disorder (SLD, Regional standardized tests used in clinical services for diagnosis of SLD-AIIMS SLD in Hindi; Mehta and Sagar, 2003 and NIMHANS index of SLD in English; Kapur et al., 1991). Magnetic resonance imaging (MRI) and functional MRI (fMRI) were carried out on 3T MR scanner (Achieva 3.0T TX, M/s. Philips Healthcare, Amsterdam, The Netherlands) with 32-channel head coil (head supported and immobilized). MRI scanning for anatomical screening was done with T1-weighted 3D sagittal images (if any incidental neurological lesion/abnormalities; parents and neurologist were informed and subject excluded from the study). In task-based fMRI response of the brain is recorded as signal modulation, as blood-oxygen-level-dependent (BOLD) changes due to the neural activity. fMRI (BOLD) cortical activity was scanned using multi-slice T2* enhanced gradient echo-echo planar imaging (GE-EPI) in axial plane with sequence parameters FOV 230mm; 31 slices and slice thickness of 5 mm without any slice gap; TR of 2 s, 163 dynamics. Visual stimulus was presented using E-prime (version 1.1, Psychology Software Tools Inc, USA) on a MR compatible LCD monitor (ESys, Invivo Corp, M/s. Philips Healthcare) with a resolution of 2560x1600@60Hz and brightness approximately 400 cd/m^2^.

### 2.1. Non-Symbolic Visual perception

Children with visual impairments were excluded from the study. The children using spectacles were assessed in adapted (comfortable) corrected vision (inside as well as outside the scanner).

#### 2.1.a. Picture Completion

This subset is used to assess ‘whole to part discrimination’ performance (MISIC subtest; Malin, 1966) that explores visual search, processing, reasoning, ability to observe details and recognize specific features of the environment by deliberately focusing attention (Wechsler, 2004a, b). The task includes 20 black line-drawings on white background where some part of the picture is missing (e.g. a girl’s face with missing lips, housefly with one antenna missing, etc) with gradually increasing picture complexity and the (missed) feature becoming miniature. The response is to search and name the missing part.

To investigate the neuro-biological aspect of visual perception, the same paradigm was designed and standardized for fMRI. Clinical assessment was done after the fMRI session (two weeks later) to avoid the priming effect. The paradigm is designed (block) as four alternate sets (ARARARAR) of active-rest blocks where ‘A-active block’ of task performance, each followed by non-active ‘R-rest block’. Each active block consisted of 4 pictures as stimuli, where the stimulus presentation time for response was standardized. The volunteers (n = 12) who participated in time standardization (outside as well as inside the scanner) were not recruited in research study. Baseline neural activity in the paradigm consisted of an initial 60 second screen display with black (30s) then white (30s) and R-rest blocks of white screen. Depending on task difficulty (standardized) stimulus presentation time in first block was 6s, in second block 10s, in third 12s and in fourth block each event was of 16s duration. Total acquisition time (including all the baselines and stimulus blocks) was 326s.

#### 2.1.b. Visuospatial Processing and Executive Control

In dyslexia research there are two opinions (Diehl et al, 2014), one as impairment in visuospatial thinking (Siok et al, 2009) and second as visuospatial strength (Brunswick et al, 2010; Wang et al, 2016). So in present study visuospatial perception is evaluated as pre-symbolic to symbolic levels (visuospatial perception, fluid reasoning and executive control), executive expression (visual discrimination, symbolic encoding for writing), perceptual processing speed (coding-digit symbols).

#### 2.1.b. Picture concept

Object Assembly is a subtest of MISIC / WAIS in performance quotient where the subject has to arrange parts of a picture puzzle in specified time, based on an underlying concept of an object (e.g. horse). It measures the ability to process nonverbal concepts quickly and accurately for categorization (skill at recognizing the conceptual common features) as an indicator of perceptual reasoning and problem solving (Malin, 1969; Wechsler, 2004a, b)

#### 2.1.b. Visuospatial Execution

Block Design subtest (of MISIC and WISC) is an indicator of perceptual reasoning and executive function (Doron et al, 2015) that measures preference to ‘learn during execution’ (mentally “putting together” complex objects by observing and manipulating its individual parts), creating solutions, and then testing them with the preferences for visual information in novel and unexpected situations (Malin, 1969; Wechsler, 2004a, b) i.e. prehension, self-doing, visuo-motor learning by experience. It includes nine red and white square blocks where the subject has to arrange these wooden cubical blocks according to the pictorial design depicted in stimulus in specified time (scored for accuracy and speed).

### 2.2. Symbolic Visual Perception

#### 2.2.i. Visual Discrimination (Pattern recognition)

It included pattern recognition of pictorial icon (<, >, c, ᴐ, ˩, ˥, etc and visual discrimination (subtest of NIMAHNS SLD battery) of two similar shapes (circle, oval, triangle), single-letter (alphabets), numbers, words (2-4 letter string), pseudosymbols where the tasks were in gradually increasing complexity.

#### 2.2.ii. Encoding non-symbolic to symbolic relation

In coding subtest of MISIC (or WISC), the performance is based on pairing number (symbolic) with an icon (non-symbolic) based on the pairs given, and (in the empty box) draw ‘an icon’ under the number (encoding and execution = decoding-encoding skill), visual-motor coordination, execution and processing speed (Wechsler, 2004b). After a few practice trials (not included in scoring), in 120 seconds the participant has to code maximum numbers-icon pairs.

#### 2.2.iii. Reading

Symbolic decoding tasks consisted of alphabet reading (upper-/ lower-case in English and in Hindi), number reading and reading at word (lexical) level in both languages.

### 2.3. Visuomotor Coordination (Symbolic execution)

Letter formation, encoding-decoding errors and visuomotor coordination was also evaluated during writing (copying) at alphabet / number, word and sentence levels (in English upper/ lower case and Hindi).

### 2.4. Auditory Perception (Phonological awareness)

In dyslexia auditory and phonological deficits are reported (Gori and Facoetti, 2015; Pennington et al, 1990; Peterson and Pennington, 2012; Snowling, 2000; Wagner and Torgesen, 1987). In present study children were excluded with hearing impairment (tested for pure-tones and speech-reception). Auditory-verbal discrimination was assessed at syllabic, lexical and minimal pairs (monosyllabic word pairs in both English and Hindi languages).

### 2.5 Data Analysis

Behavioural performance data was analyzed using SPSS version23 (IBM SPSS Statistics, 2015). Processing of fMRI data was done using Statistical Parametric Mapping package (version SPM12; Friston et al, 2007) on MATLAB (version 7.12.0.635; R2011a, MathWorks Inc., US) platform. Acquired EPI images were pre-processed involving realigning, motion correction, normalization and smoothing steps (Friston et al., 1996). At individual level (each participant) for estimation of neural activity (BOLD signal the indirect measure), postprocessing was done with general linear model, based on contrast of active to rest blocks. At second level, the group brain activity maps were estimated as BOLD clusters (1 cluster: 2x2x2 mm^3^ volume) for healthy controls (HC) and dyslexic children (DDC) with one-way ANOVA (p<0.001, cluster threshold 10) and significant activation (clusters of whole group) was overlaid on standard render template of SPM. These maps of BOLD cluster regions (coordinates) were converted from MNI to Talairach coordinates by Ginger ALE software (Turkeltaub et al, 2012; BrainMap GingerALE version 2.3, Research Imaging Institute, University of Texas Health Science Center, San Antonio) and then were classified based on Talairach atlas using Talairach Client software (Lancaster et al, 1997; Talairach and Tournoux 1988; TalairachClient 2.4.3, Research Imaging Institute, University of Texas Health Science Center, San Antonio).

## 3. Results

Sample size recruited was 75, with 50 dyslexic children (DDC) and 25 age-gender matched healthy children (HC). Out of these the data was analyzed of 67 subjects (47 dyslexics and 20 controls) excluding children with motion-artifact (head movement during fMRI acquisition greater than 2mm in translation and 2° in rotation). Both groups were compared for age, socioeconomic status, family education and bilingualism (English and Hindi; deep and shallow orthography respectively) considering bilingualism based on language acquisition-age, exposure and preference (as communication-modality in different situations, text-reading, academic-medium, media-exposure). Including mean age (HC: 12.06±2.33 years and DDC: 11.25±2.50 years) differences between the two groups for demographic variables and bilingualism were statistically non-significant. Both the groups had similar preference of Hindi for communication-mode at home, peer-group and English as academic-medium, communication-mode at school.

### 3.1. Non-symbolic visual perception

In experiment-1 (i) the visual perceptual processing was assessed with picture completion task (behavioural performance and fMRI).

#### 3.1.a. Picture Completion

Behavioural raw scores of the subtest were converted to test quotient scores (TQ) based on MISIC scaled-score table. The range of scores was suggestive of higher variability in dyslexics (DDC 51 points; HC 25 points; Table 1A, i) and performance distribution represented as most (85%) of the healthy-controls scored between 90-110 whereas 69.9% of dyslexics scored average 90-120. None of the HC scored in 120-130 or 70-90 range whereas 18.7% of DDC scored 70-90 and 11.4 % (six) of them scored between 120 and 130. Although the mean group comparison was statistically non-significant (t = -0.790; p = 0.432, two-tailed) when analyzed with independent samples t-test (Levene’s test for equality of variance f = 15.785; p < 0.0001, equal variances not assumed).

**Table 1(A).**
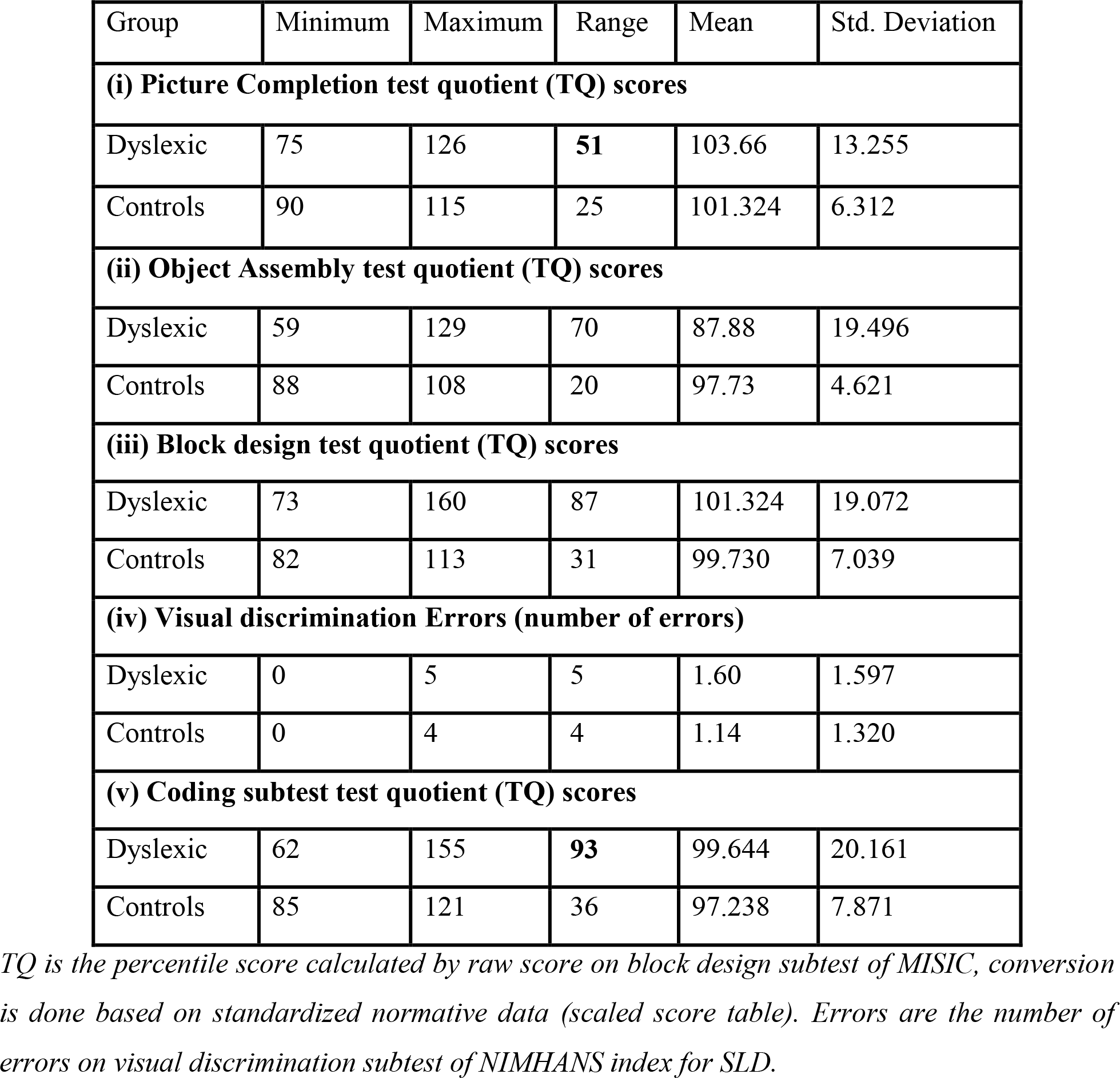
Comparing mean between dyslexic (n=47) and age-matched healthy control (n = 20) children

**Table 1(B).** Comparing mean errors between dyslexic and healthy control children for symbolic reading (dyslexic n = 47, control n = 20)

| Group |  | Mini<br>mum | Maxi<br>mum | Range | Mean | Std.<br>Deviation | Mode |
| --- | --- | --- | --- | --- | --- | --- | --- |
| Dyslexic | Lowercase English Alphabets* | 0 | 4 | 4 | 1.90 | 1.433 | 0 |
|  | Uppercase English Alphabets* | 0 | 2 | 2 | 0.08 | 0.351 | 0 |
|  | Hindi Alphabet reading<sup>+</sup> | 0 | 6 | **6** | 4.02 | 1.327 | **4** |
|  | Number reading<sup>!</sup> | 0 | 5 | 5 | 2.70 | 1.350 | **3** |
|  | English Alphabet writing Errors<sup>†</sup> | 0 | 6 | **6** | 1.04 | 1.532 | 0 |
| Controls | Lowercase English Alphabets* | 0 | 2 | 2 | 0.14 | 0.468 | 0 |
|  | Uppercase English Alphabets* | 0 | 0 | 0 | 0.00 | 0.000 | 0 |
|  | Hindi Alphabet reading<sup>+</sup> | 0 | 2 | 2 | 0.55 | 0.800 | 0 |
|  | Number reading<sup>!</sup> | 0 | 5 | 5 | 2.59 | 2.443 | 0 |
|  | English Alphabet writing Errors<sup>†</sup> | 0 | 1 | 1 | 0.05 | 0.213 | 0 |

\* number of erroneous alphabets while reading in English (lowercase / uppercase alphabets) , Alpha upp Eng- number of erroneous alphabets while reading in English, <sup>+</sup>number of errors in Hindi alphabet reading, <sup>!</sup>number of digits on which error were committed, <sup>†</sup>number of letters erroneous in form while writing English.

Brain areas of healthy children (HC) involved right hemispheric middle occipital gyri, fusiform, superior (BA 10, 6), middle (BA 8, 9), inferior frontal (BA 45) gyri, pre-central (BA 9), post-central (BA 5) gyri and left hemispheric middle, inferior occipital gyri, superior temporal (Brodmann Areas BA 22, 38), left parahippocampal gyri (BA 34) and precentral, inferior frontal (BA 9, 47) gyri. Bilateral activity was observed in precuneus and superior parietal gyri (Table 2, a; Figure 1, i).

**Figure 1.**
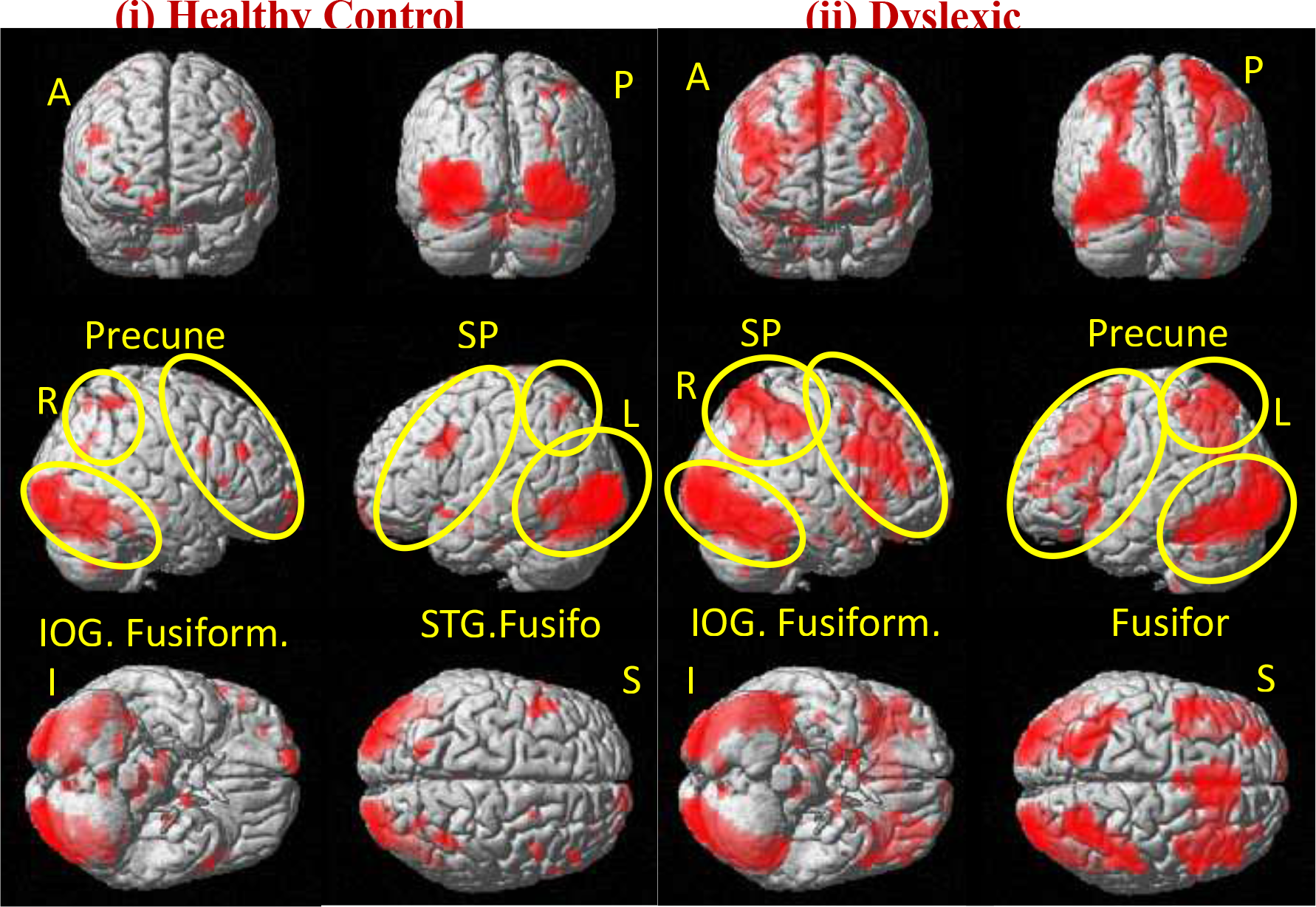
BOLD activation during visual perception task in (i) healthy control children (HC, n = 20) and (ii) dyslexic children (DDC, n = 47) (Group analysis using one-way ANOVA, p < 0.001, voxel threshold: 10); BOLD activation (important areas marked as yellow circles) overlaid on standard template render as *A-anterior view; P-posterior; R-right; L-left; I-inferior; S-superior view; meaningful word processing in (i) Healthy adults, (ii) healthy controls (iii) dyslexic children and (iv) all the three groups overlaid; IFG-inferior frontal gyrus, MFG-middle frontal gyrus, MTG-middle frontal gyrus, IOG-inferior occipital gyrus*

**Table 2(a).** BOLD activation during picture completion task in healthy control children (n=20) using one sample t-test group analysis (z score cutoff: 3.0, p < 0.001, voxel threshold: 10, 1 voxel: 2x2x2 mm^3^)

| Hemisphere | Area | BA | No. of voxels | MNI Coordinates | Z-Score |
| --- | --- | --- | --- | --- | --- |
| Right | Middle Occipital Gyrus<br>Fusiform Gyrus | BA 18<br>BA 19<br>BA 37 | 2566 | 22 -92 -4<br>26 -90 -12<br>42 -60 -16 | 6.43 |
| Left | Inferior Occipital Gyrus<br>Middle Occipital Gyrus | BA 17<br>BA 18 | 3229 | -30 -90 -6<br>-32 -92 2 | 6.31 |
| Left | Superior Temporal Gyrus | BA 22,<br>38 | 47 | -52 14 -12 | 4.13 |
| Left | Parahippocampal Gyrus | BA 34 | 28 | -14 -2 -20 | 3.77 |
| Right | Superior Parietal Lobule | BA 7 | 67 | 38 -52 58 | 4.55 |
| Right | Precuneus | BA 7 | 17 | 26 -66 56 | 3.77 |
| Right | Precuneus | BA 31, 7 | 58 | 30 -70 26<br>28 -62 36 | 3.62 |
| Left | Precuneus<br>Superior Parietal Lobule | BA 7 | 85 | -18 -66 58<br>-22 -58 52 | 4.23 |
| Right | Superior Frontal Gyrus | BA 10 | 27 | 30 66 -2 | 3.80 |
| Right | Superior Frontal Gyrus | BA 6 | 12 | 22 10 72 | 3.37 |
| Right | Superior Frontal Gyrus | BA 10 | 94 | 6 68 -12 | 3.67 |
| Right | Middle Frontal Gyrus | BA 11 | 10 | 32 48 -22 | 3.66 |
| Right | Middle Frontal Gyrus | BA 8,9 | 55 | 54 34 34 | 3.62 |
| Right | Precentral Gyrus | BA 9 | 48 | 46 10 32 | 3.63 |
| Right | Postcentral Gyrus | BA 5 | 15 | 38 -38 80 | 3.62 |
| Right | Inferior Frontal Gyrus | BA 45 | 12 | 58 22 8 | 3.58 |
| Left | Precentral Gyrus<br>Inferior Frontal Gyrus<br>Middle Frontal Gyrus | BA 6<br>BA 9 | 224 | -44 10 36<br>-48 22 28<br>-46 12 24 | 4.26 |
| Left | Inferior Frontal Gyrus | BA 47 | 24 | -60 20 -14 | 3.19 |
| Left | Lentiform Nucleus<br>Lateral Globus Pallidus |  | 14 | -24 -14 -10 | 3.56 |
| Right | Substantia Nigra<br>Brainstem |  | 56 | 12 -20 -10 | 3.89 |
| Right | Cerebellar Tonsil |  | 41 | 32 -62 -46 | 4.49 |
| Right | Cerebellar Tonsil |  | 32 | 18 -46 -44 | 3.58 |
| Left | Cerebellar Culmen |  | 77 | -6 -34 -4 | 4.42 |
| Left | Cerebellar Culmen |  | 15 | -36 -44 -28 | 3.82 |
| Left | Cerebellar Nodule |  | 69 | -6 -54 -32 | 4.20 |
| Left | Cerebellar Pyramis<br>Declive |  | 32 | -6 -80 -26<br>-2 -74 -22 | 3.51 |

BA: Brodmann Area, MNI: Montreal Neurological Institute

In dyslexic children, activity was observed in right hemispheric inferior, middle occipital, superior parietal, inferior parietal gyri, hippocampal, parahippocampal gyri, superior frontal gyrus (BA 6, 8, 9), inferior frontal gyrus (BA 11) and left hemispheric precuneus, superior temporal gyri, amygdala, middle frontal (BA 9, 10), superior frontal, orbital frontal, precentral gyri with along with bilateral fusiform gyri, superior, middle, medial (left BA 10, right BA 25) and inferior frontal gyri (left BA 47, right BA 46) (Table 2,b; Figure 1, ii).

**Table 2(b).** BOLD activation during picture completion task in dyslexic children (n=47, Group analysis using one sample t-test, p < 0.001, voxel threshold: 10)

| Hemisphere | Area | BA | No. of voxels | MNI Coordinates | Z-Score |
| --- | --- | --- | --- | --- | --- |
| Right | Inferior Occipital Gyrus<br>Middle Occipital Gyrus<br>Fusiform Gyrus | BA 18<br>BA 37 | 4935 | 28 -92 -4<br>36 -86 -8<br>44 -62 -16 | 7.53 |
| Right | Fusiform Gyrus | BA 20 | 12 | 40 -14 -30 | 3.76 |
| Left | Fusiform Gyrus | BA 19<br>BA 18 | 3901 | -26 -90 -12<br>-20 -96 -8 | 7.36 |
| Left | Superior Temporal Gyrus | BA 38 | 39 | -26 12 -58 | 3.63 |
| Right | Superior Parietal Lobule<br>Inferior Parietal Lobule | BA 7<br>BA 40 | 2132 | 38 -54 56<br>46 -32 44 | 6.02 |
| Left | Precuneus | BA 7 | 1555 | -24 -64 44 | 4.69 |
| Right | Superior Frontal Gyrus<br>Precentral Gyrus | BA 8<br>BA 6<br>BA 9 | 4404 | 4 32 50<br>4 22 54<br>50 12 32 | 5.97 |
| Right | Middle Frontal Gyrus | BA 6 | 37 | 38 2 52 | 3.85 |
| Right | Medial Frontal Gyrus | BA 25 | 42 | 14 8 -28 | 3.71 |
| Right | Superior Frontal Gyrus | BA 10 | 19 | 18 76 -16 | 3.39 |
| Right | Inferior Frontal Gyrus | BA 11 | 141 | 32 30 -36 | 4.21 |
| Left | Middle Frontal Gyrus<br>Precentral Gyrus | BA 9<br>BA 10<br>BA 6 | 1526 | -46 14 38<br>-32 50 14<br>-34 6 30 | 5.79 |
| Left | Superior Frontal Gyrus | BA 6 | 97 | -2 2 68 | 4.62 |
| Left | Middle Frontal Gyrus | BA 6 | 15 | -30 2 66 | 3.47 |
| Left | Superior Frontal Gyrus | BA 10 | 14 | -4 66 -22 | 3.33 |
| Left | Middle Frontal Gyrus | BA 10 | 37 | -26 74 -4 | 4.29 |
| Left | Medial Frontal Gyrus | BA 10 | 24 | -16 68 -14 | 3.75 |
| Left | Middle Frontal Gyrus | BA 11 | 90 | -22 32 -26 | 3.75 |
| Left | Inferior Frontal Gyrus | BA 46 | 15 | -50 42 -8 | 3.78 |
| Left | Inferior Frontal Gyrus | BA 47 | 79 | -52 16 -10 | 4.53 |
| Left | Orbital Gyrus | BA 47 | 80 | -26 26 -40 | 4.32 |
| Left | Insula<br>Clastrum |  | 114 | -42 16 -4<br>-30 22 2 | 3.85 |
| Right | Hippocampus<br>Thalamus |  | 42 | 32 -32 -2<br>24 -28 -4 | 4.03 |
| Right | Parahippocampal Gyrus<br>Lentiform Nucleus | BA 34<br>MGP | 113 | 22 0 -22<br>16 0 -14 | 4.72 |
| Left | Parahippocampal Gyrus (Amygdala)<br>Medial Globus Pallidus |  | 30 | -20 -4 -14<br>-12 -4 -16 | 4.50 |
| Right | Substantia Nigra, Brainstem |  | 33 | 14 -22 -10 | 4.17 |
| Right | Cerebellar Tonsil |  | 16 | 32 -64 -50 | 4.33 |
| Right | Cerebellar Pyramis |  | 48 | 8 -80 -24 | 4.19 |
| Right | Caudate Body |  | 38 | 20 10 10 | 3.93 |
| Left | Cerebellar Culmen |  | 55 | -8 -30 -14 | 4.18 |
| Left | Cerebellar Uvula |  | 39 | -6 -78 -28 | 3.95 |

Group comparison of DDC and HC (DDC > HC), revealed significant activation clusters in the left hemispheric superior temporal gyri (BA 20), inferior parietal lobule (BA 40), orbital frontal (BA 47, 11), left uncus (BA 20) and anterior cingulate in limbic lobe. BOLD activity was also seen in right hemispheric inferior (BA 47), middle (BA 8, 9) and medial (BA 32) frontal gyri (Table 2,c). Comparison of HC group with DDC (HC > DDC), significant activation regions were left hemispheric middle, inferior occipital gyri (BA 18) and cuneus with few clusters in right hemispheric middle frontal gyrus (Table 2,c).

**Table 2(c).** 4. Comparing BOLD activation during picture completion task in healthy control (n=20) and dyslexic (n = 47) children (Group analysis using one sample t-test, p < 0.001, voxel threshold: 10, 1 voxel: 2x2x2 mm^3^)

| Hemi-sphere | Area | BA | No. of voxels | MNI Coordinates | Z-Score |
| --- | --- | --- | --- | --- | --- |
| **Healthy control versus dyslexic children** |  |  |  |  |  |
| Left | Cerebellar Nodule |  | 16 | -8 -54 -32 | 3.85 |
| Left | Cerebellar Culmen |  | 18 | -6 -36 -4 | 3.71 |
| Left | Middle Occipital Gyrus<br>Inferior Occipital Gyrus | BA 18 | 113 | -30 -90 2<br>-38 -96 -8 | 3.58 |
| Right | Middle Frontal Gyrus | BA 8 | 12 | 54 34 34 | 3.53 |
| Left | Cuneus | BA 18 | 22 | -20 -86 34 | 3.39 |
| **Dyslexic versus healthy control children** |  |  |  |  |  |
| Left | Orbital Gyrus | BA 11 | 51 | -14 42 -46 | 3.16 |
| Left | Clastrum |  | 52 | -32 10 4 | 4.10 |
| Left | Orbital Gyrus | BA 47 | 12 | -16 22 -40 | 4.02 |
| Left | Uncus | BA 20 | 35 | -38 -14 -32 | 3.98 |
| Left | Lentiform Nucleus | Putamen | 25 | -22 12 -14 | 3.82 |
| Right | Inferior Frontal Gyrus | BA 47 | 50 | 54 44 -32 | 3.81 |
| Left | Anterior Cingulate | BA 24 | 16 | -8 36 -8 | 3.59 |
| Right | Middle Frontal Gyrus | BA 8,9 | 88 | 30 32 34 | 3.58 |
| Left | Inferior Parietal Lobule | BA 40 | 15 | -62 -30 24 | 3.38 |
| Right | Lentiform Nucleus | Putamen | 21 | 22 10 -10 | 3.33 |
| Right | Medial Frontal Gyrus | BA 32 | 15 | 20 20 42 | 3.23 |

#### 3.1.b. Visuospatial Reasoning and Executive Control

Experiment-1 part (b) consisted of behavioural performances on picture-concept visuospatial reasoning (object assembly) and executive function (block design sub-test)

#### 3.1.b. Picture concept

Object Assembly mean (DDC-87.88, HC-97.73) and median (DDC-86.00, HC-99.50) scores were distinguishable whereas mode was 100 in both the groups (table 1A,ii). Distribution of scores most (90%) of the HC performed average (90 to 110) whereas in dyslexic scores ranged differently (59 to 129). Variability in distribution of scores as 40.7% of DDC performed slower (60-70 in 21.6%, 70-80 in 19.1%), 14.4% borderline (80-90), average performance in 26.3% (90-100 score in 19.1%, 100-110 in 7.2%) and 16.8% performed faster (110-120 in 9.6%, 120-130 in 7.2% dyslexics). Mean scores were significant different in two groups (t = 2.327; p = 0.023 i.e. p < 0.05, two tailed; Levene’s test f = 37.421; p < 0.001, equality not assumed).

#### 3.1.b. Visuospatial Execution

Performance for Block-design subtest (table 1A,iii) and distribution of scores in dyslexics show variability. Majority (85%) of HC and 64.8% of the DDC scored between 90-110. Performers variability in dyslexics with 13.5% below average (slow, 70-80 TQ), 5.4% borderline (80-90) and 18.9% scored above average (2.7% scored 110-120, 8.1% 120-130, 8.1% 130-160). On two-sample ‘t’test, scores were not significantly different (t = -0.377; p = 0.708 two tailed equal variance not assumed).

### 3.2. Symbolic Visual Perception

Experiment-2 investigated nonsymbolic to symbolic interface with pattern recognition, coding and reading performances.

#### 3.2.i. Visual Discrimination (Pattern Recognition)

Mean scores for symbolic visual discrimination (as number of errors committed) were not significantly different in two groups (t = -1.173, p = 0.245, two tailed; Levene’s equality; f = 2.172, p = 0.145) (table 1A,iv). In both groups ‘no-errors’ were observed for shapes, single-symbol, alphabet or two-alphabet words (e.g. HE). Difference in dyslexic and healthy readers were observed at 3-5 alphabet/string words (SHIP, SLIP) and number of errors committed (3-erronous alphabets DCC-17%, HC-5% and 5-erronous alphabets DCC-6.4%, HC-0%).

#### 3.2.ii. Encoding non-symbolic to symbolic relation

Mean difference in two groups for coding subtest was non-significant (with two-sample t-test 0.535; p = 0.594, two tailed), but dissimilarity was seen in correct-encoding score (median HC-100; DCC 97) and frequently occurring score (mode HC-99; DCC-90) (table 1A, v). Distribution of scores showed variability in DCC (HC 95% scored 90 to 120; DCC range was 62 to 155 with 69.2% scoring 80-110). Variability in dyslexic group observed as slow performers (4.2% scored 60-70, 12% scored 70-80) and fast performers (7.2% scored 110-120, 4.8% ranged 140-150, and 2.4% scored 150-160). Differences were not due to developmental motor incoordination as (motor) history estimated with Mann-Whitney-U test (Z= -0.951; p = 0.342, two-tailed) was statistically non-significant, though 10% of HC and 21.3% of DDC had positive history of motor delay.

#### 3.2.iii. Reading (Alphabet and Number)

The symbol identification skill was assessed by performance on reading single alphabet in English (upper- and lower-case alphabets), Hindi (varnama:la: or alphabets) and number reading at one, two, three, four or five -digit level. The distribution of errors (Figure 2,i) and mean alphabet reading in English (lowercase) was significantly different (t-test t = -5.594, p < 0.0001, two tailed; Levene’s test, f = 31.062, p < 0.0001. English uppercase alphabet reading the distribution of errors (no errors in 100% HC and 93.6% dyslexics) and mean score were not significantly different (f = 5.708, p = 0.02; t = -1.134, p = 0.261, 2-tailed). In Hindi alphabet reading mean errors were significantly different in two groups (f = 1.183, p = 0.281; t = -11.334, p < 0.0001, two-tailed) and four alphabets were frequently (mode) read erroneously in dyslexic group. Mean digit (number) reading errors in HC and DDC groups were not significantly different (f = 48.131, p < 0.0001, equality of variance not assumed; t = -0.244, p = 0.808, 2-tailed) (table 1B). The reading errors at lexical level (meaningful word-lexical level) were related to the vowel duration or consonant clusters (phonological) (Figure 2,ii). Number string reading in HC group no-errors were seen in 45%, 4-digit numbers erroneous in 15% (3) and 5-digit erroneous in 40%. In DDC group, no-errors in 14.9%, 2-digit number erroneous in 17%, 3-digit in 38.3%, 4-digit number in 27.7% and 2.1% (one dyslexic) had error on 5-digit numbers that means 55.3% of dyslexic children read erroneously 2- and 3-digit numbers onwards.

**Figure 2(i).**
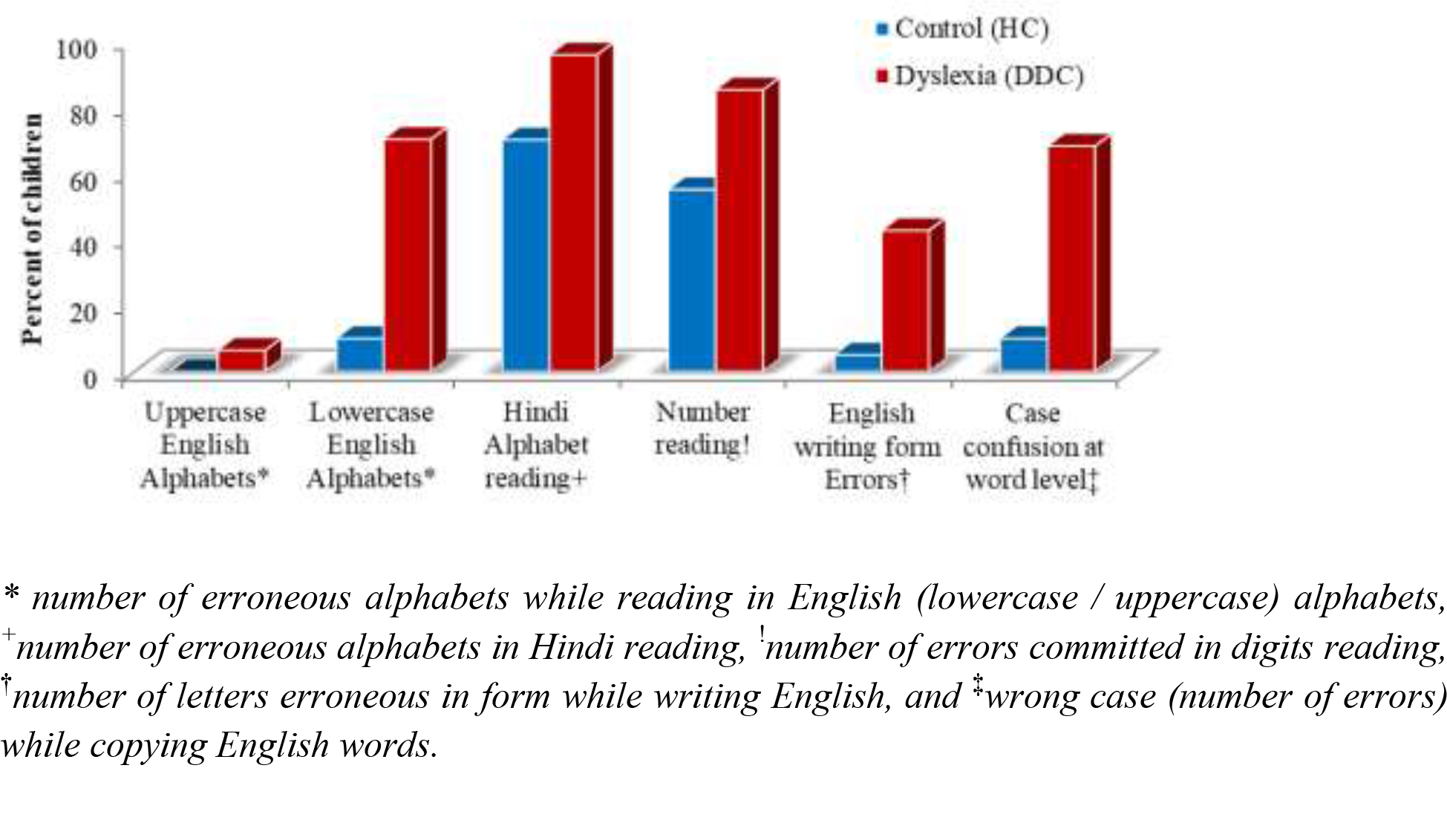
Comparing dyslexic (n = 47) and healthy (n = 20) children for symbolic performance (percent of children having errors)

**Figure 2(ii).**
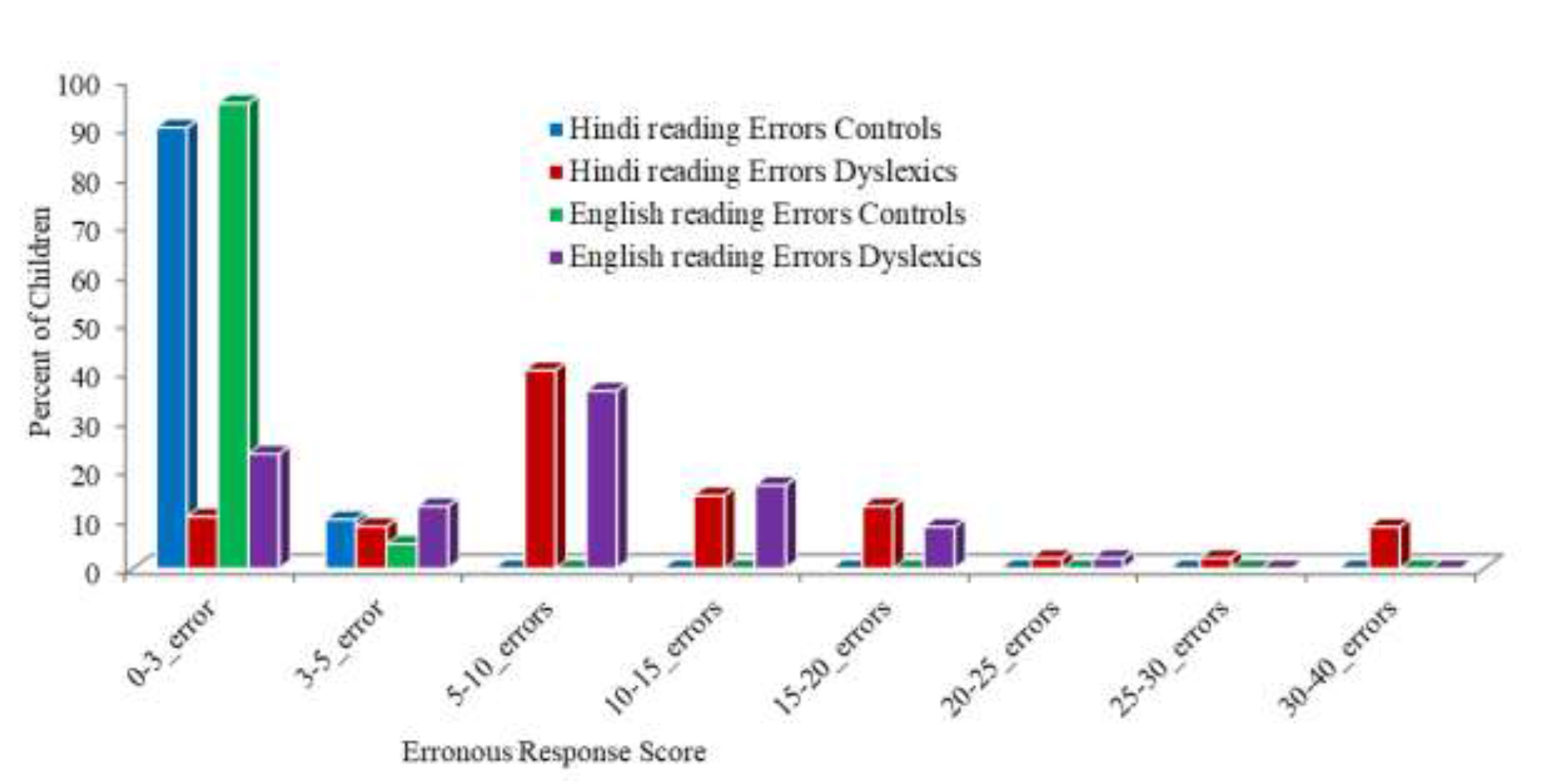
Comparing Reading Errors in dyslexic (DDC, n = 47) and healthy (HC, n = 20) for two languages

**Figure 2(iii).**
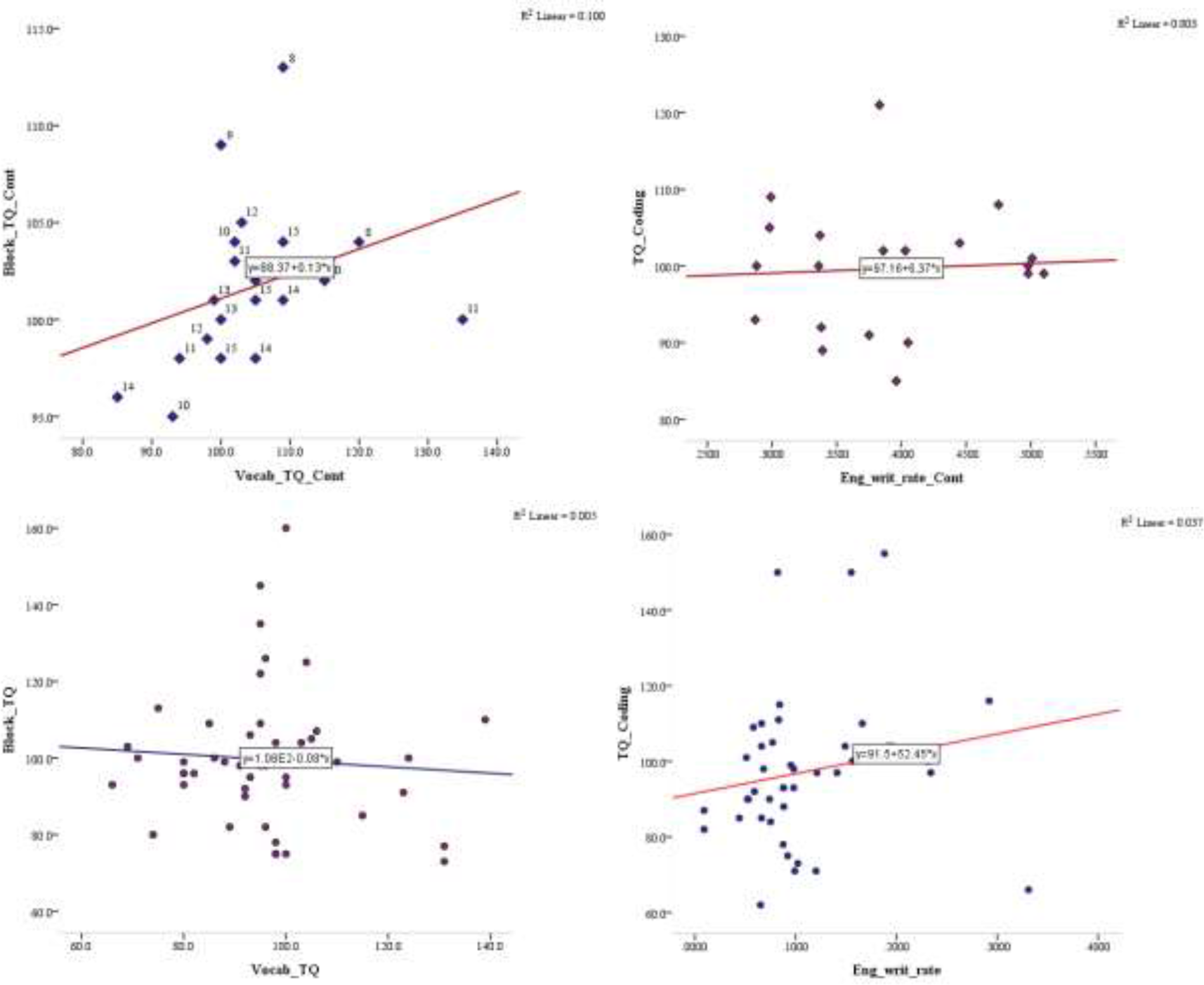
Association of nonsymbolic to symbolic execution (a) Block-design with vocabulary; (b) Encoding speed with English writing rate (while copying) in typical readers (HC, n = 20 and DDC n = 47) (scatter-plots) [*Block_TQ_Cont are scores of HC for Block-design subtest and Block_TQ are scores of DDC; Vocab_TQ_Cont means Vocabulary scores of HC and Vocab_TQ means vocabulary scores DDC; Coding_TQ are the scores of Coding subset; Eng_writ_rate_Cont means English writing rate of HC while copying; Eng_writ_rate means English writing rate of DDC while copying*]

**Figure 2 (iv).**
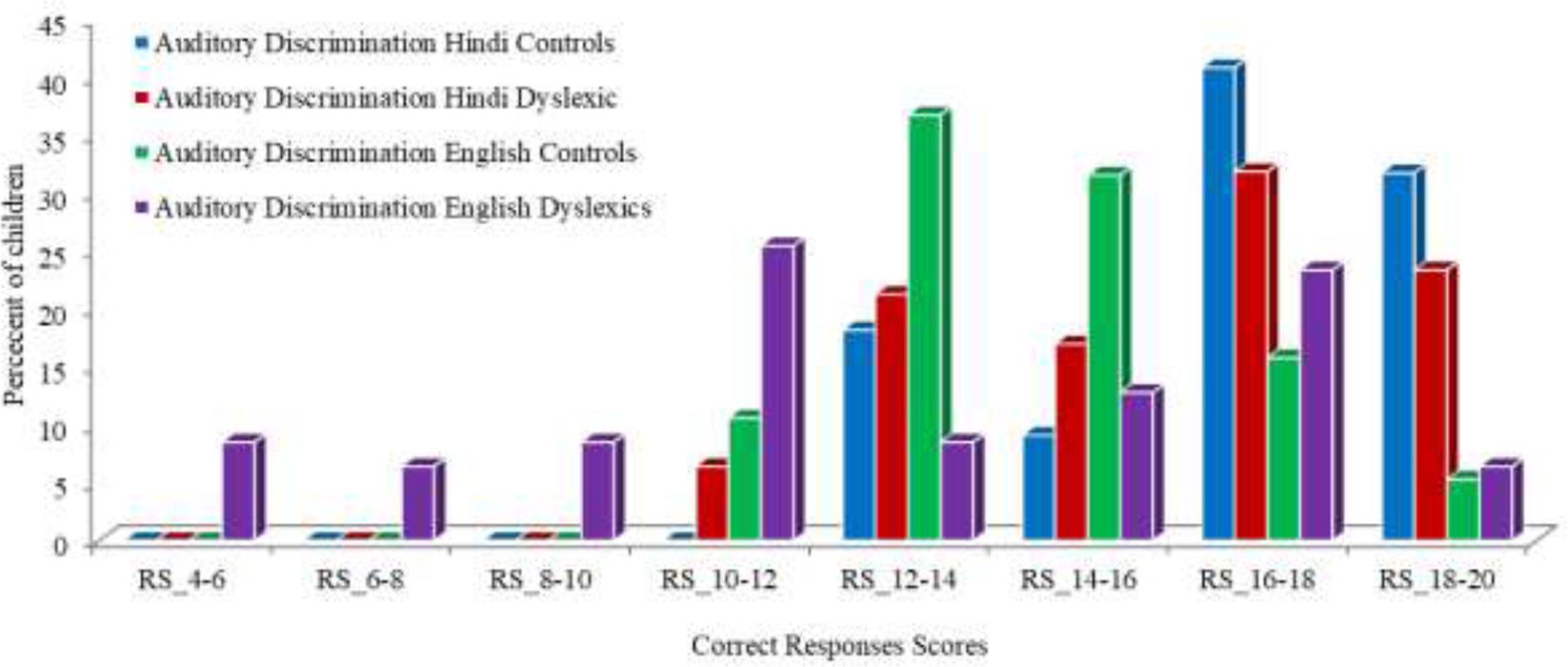
Comparison Phonological Awareness between dyslexic (DDC, n = 47) and healthy control (HC, n = 20) children for two languages at word level

**Figure 2(v).**
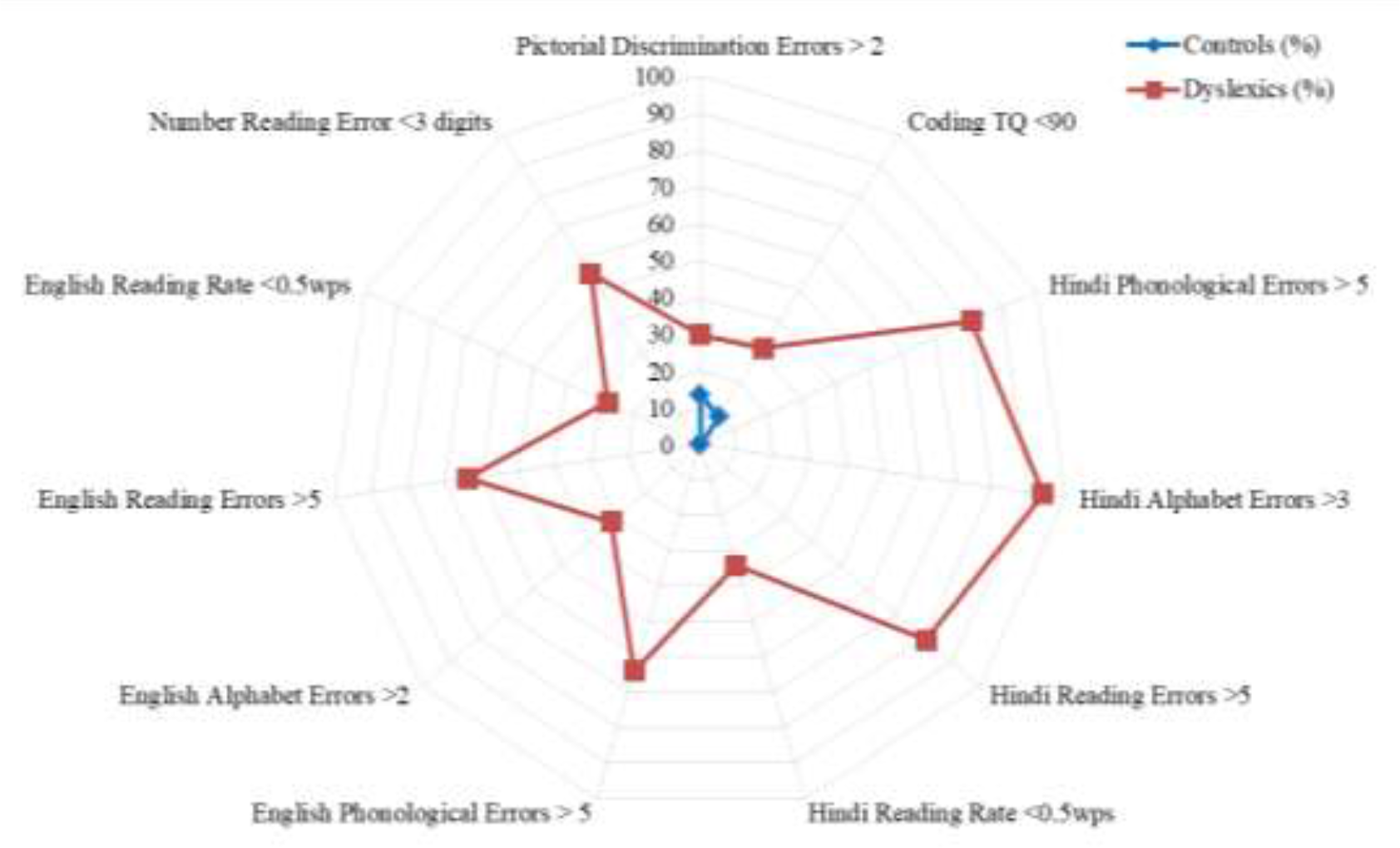
Comparing dyslexic (DDC, n = 47) and healthy (HC, n = 20) children nonsymbolic (pictorial) to symbolic (text at single alphabet and string level) performance; *(number of children represented in %)*

### 3.3. Symbolic Visuomotor Execution

Experiment-3 evaluated symbolic coordination and execution as letter formation in writing (copying). Errors in English copying were significantly different (t = -3.028, p = 0.003, two-tailed; f = 31.300, p < 0.0001, equality of variance not assumed). Differences in distribution of errors were also seen as in HC 95% had no letter-form errors, 5% had one error perseveration in ‘m’ with extra arm. Whereas in DCC 57.4% had no-errors, 10.6% had single-alphabet-erroneous, 17% had two-alphabets-erroneous, 4.3% 5-alphabets and 2.1% had 6-alphabets erroneous. The errors observed in DCC were mirroring (especially for c, e, s, E), perseveration (for m, n, w), vertical flip (for y as h, p as b), and irregular pattern errors (k/ l, l / t, o / a, s /s c, etc).

### 3.4. Auditory perception (Phonological awareness)

In experiment-4 comparing the two groups, significant difference in English minimal-pair auditory discrimination (t = 1.861, p < 0.05; f = 8.411, p = 0.005, equality of variance not assumed) as well as in scores’ distribution (figure 2,iv). Auditory discrimination in Hindi was non-differentiable in two groups (t = 1.631, p = 0.108; F = 2.410, p = 0.125, two-tailed) and neither in score distribution.

### 3.5. Association of nonsymbolic to symbolic performance

Association among the variables of non-symbolic and symbolic performance was analyzed with Scatter plot, Pearson’s correlation coefficient and covariance, along with exploring the monotonic relation with Spearman’s coefficient, since the variables are TQ scores (instead of raw scores) (Moore et al, 2013). In typical-reading children (HC), moderate positive correlation was found between encoding nonsymbolic to symbolic processing speed and vocabulary (coding subtest ‘r’ = 0.528, p = 0.017. The positive covariance of coding scores and vocabulary (45.779) was detected. Mazes TQ (spatial-reasoning and execution) scores were positively (weak) correlated with vocabulary but could not reach statistical significance (Correlation coefficient ‘r’ 0.409, p = 0.073). In HC negative correlation of mazes scores with writing errors (r = -0.049, p = 0.026; negative covariance of -100.687) and mazes scores covariance with erroneous alphabets (-2.150), vocabulary (62.600). Performance of block-design was also associated with vocabulary (scatter plot and monotonic relation Spearman’s ‘rho’ = 0.518, p = 0.019 in HC group but was not observed as significant Pearson’s correlation coefficient (Figure 2,iii). In dyslexic children (DDC) no such correlation was observed with spatial-reasoning and symbolic variables. Encoding speed in DDC had weak negative correlation with copying spelling-errors (r = -0.261, p = 0.076; Spearman’s ‘rho’ = -0.325, p = 0.026) negative covariance (-123.707) and weak negative correlation of block-design with erroneous alphabet scores (‘r’ = -0.283, p = 0.054; covariance = -7.230). Dyslexic children had weak significant correlation between block design and picture completion TQ scores (Pearson ‘r’ = 0.378, p = 0.009; covariance 84.676 and Spearman’s ‘rho’ = 0.455, p = 0.001.

## 4. Discussion

Reading requires integrated multifaceted cognitive processing (Giovagnoli et al, 2016) and dyslexia may be attributed to deficits of visual perception, attention, distracters’ inhibition and/or auditory perception (Ahissar, 2007; Goswami et al, 2011; Lobier et al, 2011; Menghini et al., 2010; Peterson and Pennington, 2012; Ramus and Ahisaar, 2012; Reid, 2016; Shaywitz and Shaywitz, 2008; Vidyasagar and Pammer, 2010; Zoccolotti et al, 2016). Thus in present study, to untangle the role of perception and attention in reading mechanism, assessments included non-symbolic to symbolic skills. Non-symbolic are based on non-verbal fluid reasoning (picture completion, object assembly and block design) that measure the ability of visual search, attentional focus, visuospatial, visual-motor problem solving and executive skills. The symbolic (linguistic/verbal) performances include pattern recognition, visual discrimination, alphabet/number reading, word (lexical) reading, writing (alphabets and word copying), symbolic-nonsymbolic encoding, auditory discrimination (phonological awareness) and vocabulary development.

Visual search paradigm (like picture completion task in the study) has been useful for investigating brain areas for allocation of attention, spatial localization, identification of target, and distractors inhibition (Ellison et al, 2014). Behavioural performance in DDC had higher variability (Bosse et al, 2007) but non-significant group mean difference between HC and DDC. In HC task based BOLD activation of right hemispheric fusiform (BA 19: visual association area occipital area, BA 37: temporal area) and left inferior occipital gyri (BA 17, primary visual area) with bilateral middle occipital (BA 18), attribute to visual-perception and reasoning (Ardila et al, 2015; Ellison et al, 2007; Knauff et al, 2002). Visual area V4 (BA 18, 19) of the ventral stream play a primary role in figure-ground segmentation of visual scene by attentional filtering (Roe et al, 2012). Visual search task invoked activity of right fusiform, middle occipital, superior parietal lobule, precentral, inferior frontal and left middle frontal gyri in HC (Ellison et al, 2014). Right BA 31 (precuneus) has role in visuospatial imagery, self-processing operations (first-person perspective and/or experience of agency) and episodic memory retrieval (Cavanna and Trimble, 2006). Right superior parietal lobule (BA 7) is involved in visuo-verbal interface of working memory whereas right fusiform with left precuneus (BA 7) in visual attention, spatial orientation (Ardila et al, 2015) and top-down attentional control (Friedman-Hill et al, 2003). Activation in left language areas involves visual naming (naming the missing part in picture). Left parahippocampal gyrus (BA 34) activation supports memory processing (Grasby et al, 1992; Svoboda and Levine, 2009). Activation of BA 9 with BA 44, 45, precentral (in frontal) precuneus, middle temporal, hippocampus and amygdala constitute the conflict-resolution network (Margolis et al, 2018).

The visual perception activates bilateral posterior areas while visual naming is associated with left lateralization (Ardila et al, 2015; Ellison et al, 2014; Hamberger and Seidel, 2009). Group comparison of HC to dyslexic (HC > DDC), BOLD activity in left hemispheric middle, inferior occipital gyri (BA 18) and left cuneus with few clusters in right hemispheric middle frontal gyrus attribute visuo-phonological loop processing for semantic objects naming (Baddeley, 2012; De Benedictis et al, 2014).

BOLD activation of dyslexic group was similar to that of healthy children in right inferior occipital (BA 18) and fusiform gyri (BA 37), but it was being complemented by left fusiform (BA 18, 19). The volume of BOLD activation (number of clusters 6620 in DDC, 506 clusters in HC) in frontal and parietal areas (dyslexics 3691 clusters, controls 242 clusters) was higher in dyslexic group as compared to HC. The volume of activation in precuneus (BA 7) for dyslexics was higher bilaterally (dyslexics right hemispheric 2132 clusters, 1555 left; controls right hemispheric 84 cluster, left 85) indicating an extra effort for visual attention, spatial orientation and top-down attentional control (visuospatial model; figure 3). The occipital striate (BA 17), extra-striate (BA 18, 19) with fusiform (BA 37) as ventral stream system are involved in the visual perceptual processing, figure-ground filtering and auditory linguistic function (Ardila et al, 2015; Milner and Goodale, 2008; Roe et al, 2012). On group comparison of dyslexic with control children (DDC > HC), significant activity in left hemispheric orbito-frontal gyrus, inferior parietal lobule, right hemispheric inferior, middle and medial frontal gyri was observed. These differences emphasize greater involvement of frontal and prefrontal regions in dyslexic children for visual processing and modified top-down feedback to extra-striate visual areas that play an important role in reading (Al Dahhan et al, 2020; Ardila et al, 2015; Brem et al, 2020; Friedman-Hill et al, 2003; Nash et al, 2017; Roach and Hogben, 2007; Whaley et al, 2016). Higher BOLD activity of left anterior cingulate in dyslexic group suggests conflict in feedback loop (Posner and DiGirolamo, 1998), where the decreased activity (interface) of right frontal cortex and anterior cingulate contribute to optimal attentional vigilance and conflict resolution (Margolis et al, 2018; Parasuraman, Warm, See, 1998) (visuospatial model; figure 3).

**Figure 3.**
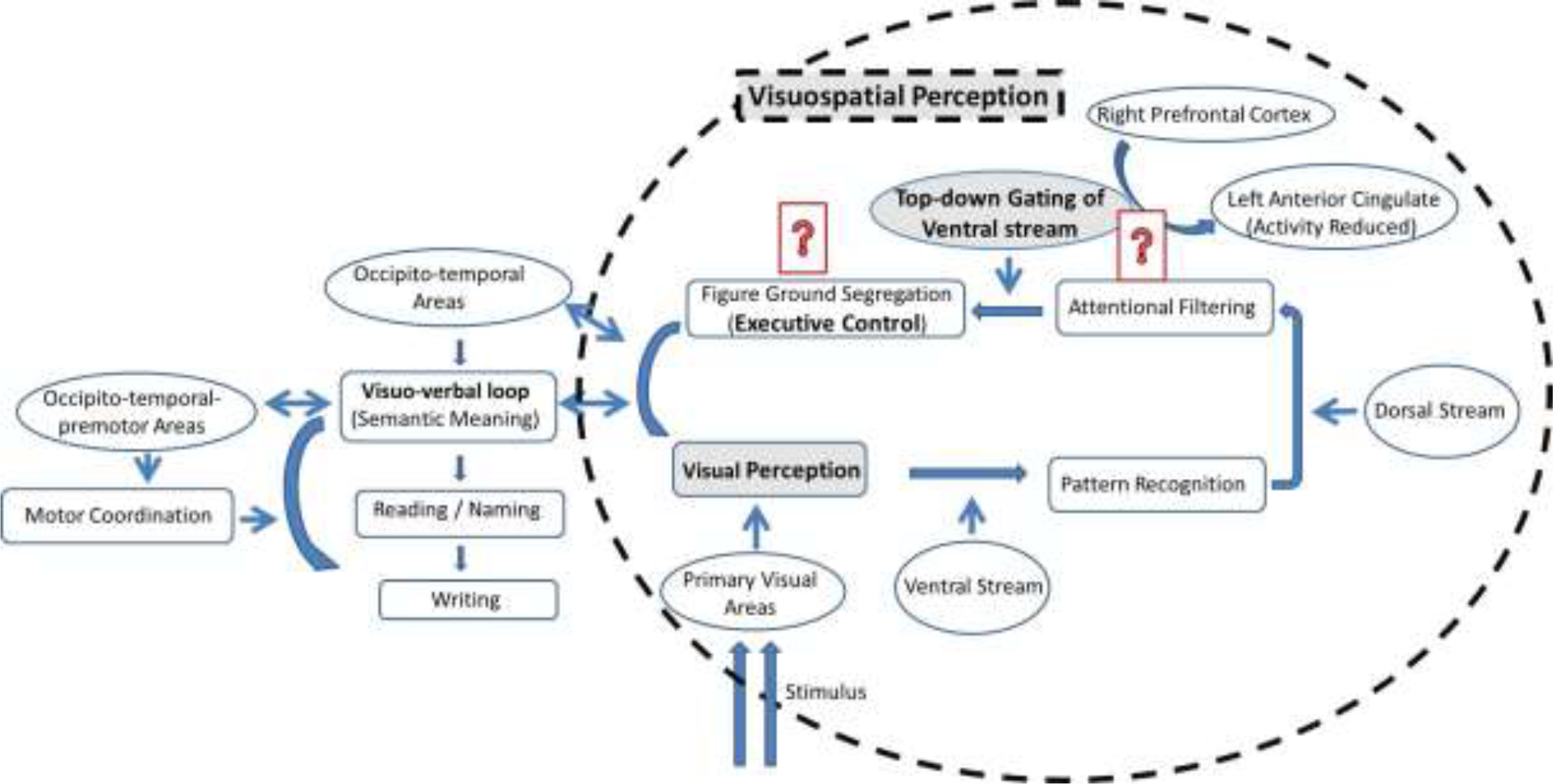
Role of Perception, attention and executive-control in Reading *(Simplified schematic diagram where two question marks represent the observed dyslexia differences at presymbolic level)*

Object assembly task is time bound visually guided actions of picture concepts (whole into part or parts into whole) involving coordination of spatial attention, localization, movement, depth perception and objects’ context dependent movement manipulation. Similar to picture completion task dyslexics distribution of scores show variability from slower (in 55.1% DDC) to faster (in 16.4%) performance rate (HC range of 90 to 120). As observed in picture completion task higher BOLD activity in ventral stream including occipital areas (BA 18 and BA 19) and fusiform gyri (occipital and temporal) attribute greater effort in visual attention and suppression of distractor stimuli (Al Dahhan et al, 2020; Pinsk et al., 2004; Roe et al, 2012) that is attentional filtering by stimulus prioritization, shrinking receptive field and noise reduction (Kastner, 2004). Deficits of attentional filtering (with unimpaired orientation and discrimination judgements) in dyslexia is not a general feature (Roach and Hogben, 2007) but is observed as performance variability (Table 1A) accompanied ventral fronto-parietal network lateralized more towards the right hemisphere (right hemispheric 2132 clusters, 1555 left) suggests modified top-down feedback (Corbetta and Shulman, 2002; Friedman-Hill et al, 2003; Kastner, 2004). This modified feedback gating to the ventral stream (primary visual cortex) may explain the performance differences (Hebart and Hesselman, 2012; Milner and Goodale, 2008; Goodale and Milner 1992; Vidyasagar and Pammer, 2010). The differences of anterior cingulate (BOLD) activity in HC > DCC and DCC > HC (group comparisons) also attribute to modified feedback loop for distractor suppression (Parasuraman, Warm, See, 1998; Posner and DiGirolamo, 1998).

In Experiment-2 visuospatial executive (block-design subtest) performance of dyslexics showed higher variability and no statistical mean difference between the two groups (Hawelka and Wimmer, 2008). In DDC group 35.1% either took longer (18.9% of dyslexics) or shorter time (16.2% of dyslexics) compared to HC. It might be hypothesized as attention driven visuospatial control for manual skill/ prehension, which is unifying function of left precuneus (BA 7; integration for visuospatial imagery) and right precuneus (BA 31; experiential self-processing spatial operations) (Cavanna and Trimble, 2006). This parietal top-down signal to visual areas coordinated with thalamus guides motor actions (Goodale and Milner, 1992; Kastner and Pinsk, 2004; Norman and Shallice, 1980; Parasuraman et al, 1998; Vossel et al, 2014). Weak correlation between picture completion and block-design subtest scores with such neural processing of attentional control, targeted selection, suppression of distractors, computational feedback in dyslexic children (figure 3) (Kastner and Pinsk, 2004; Vossel et al, 2014) further emphasize alterations at nonsymbolic level (Arrington et al, 2019; Ding et al, 2016; Giovagnoli et al, 2016).

These non-symbolic visual top-down feedback differences in dyslexic children indicate affected push-pull mechanism of visual attention (Kastner and Pinsk, 2004; Pinsk et al, 2004; Roach and Hogben, 2007; Xia et al, 2016). At symbolic level, this push-pull mechanism might affect the symbol string or letter interpretation (connectionist multi-trace model, Ans et al., 1998; Bosse et al, 2007). In present study, mean visual discrimination scores (while reading at alphabet or word level) (Shovman and Ahissar, 2006) and the distribution of scores (Hawelka and Wimmer, 2008) of two groups were not different statistically for English uppercase. Reading errors in lowercase English and Hindi, significantly increased at bigger string (3-4 alphabets level) in dyslexic children (Table 1B; Figure 2,iv). In Hindi alphabets, minor pattern differences (addition or omission of extra arm in consonant / vowel) result in major phonemic or meaning (morphophonemic) representation (Nag, 2011; Zhou et al, 2014). Such discrimination is hypothesized as “a bidirectional association” between pure visual and reading skills (Nash et al, 2017; McBride-Chang et al, 2011). Modified BOLD activation in visual search task might support where DDC had bilateral occipito-temporal fusiform (BA 37) (Hirshorn, Li, Ward, Richardson, Fiez, Ghuman, 2016), left insula, left superior temporal gyrus, left inferior and orbital frontal gyri that constitute the ventral stream for perception, recognition, discrimination, identification (Hebart and Hesselmann, 2012; Nash et al, 2017) and BA 37 (fusiform gyrus) involved in semantic processing of oral/ written language (Al Dahhan et al, 2020; Ardila et al, 2015; Cloutman et al, 2009; Nash et al, 2017; Whaley et al, 2016). These modifications might be extrapolated to higher symbolic visual discrimination errors in dyslexia (Cohen, 2016; Hebart and Hesselmann, 2012; Vidyasagar and Pammer, 2010; Vossel et al, 2014; Whaley et al, 2016) especially at bigger string level (3-4 letter words) (Hirshorn et al, 2016; Pammer, 2014) as the attention deficits (by SERIOL model- Whitney, 2001; serial allocation of attention-Vidyasagar and Pammer, 2010). In number reading most (85.1%) of the DCC were erroneous from 2 digits onwards further indicated that combination of symbol string and pattern discrimination lead to greater difficulty in allocation of visual attention (Bosse et al, 2007; Hawelka and Wimmer, 2008; Lassus-Sangosse et al, 2008).

Coding subtest measures synchronized visual discrimination, spatial control, symbolic decoding-encoding, motor execution and processing speed (Wechsler, 2004b). The mean performance of two groups was not significantly different but dyslexic children (45.5% scored average 90-120) scored differently to healthy controls (94.8%). It correlates with reading / writing ability and may be explained with differences in selective-attention, inhibitory-execution, cognitive-flexibility (visual-motor processing speed) and working-memory (complexity of coding). (Bogon et al, 2014; Easson et al, 2020; Engel de Abreu et al, 2014; Scordella et al, 2015). Analysis of writing skill (while copying words in English) evidenced statistically significant errors in dyslexics where 68.1% of DDC and only 10% (two) of the HC showed case confusion (upper/lower). In dyslexic children, errors while copying (writing) also included mirroring, perseveration, vertical flip or irregular forms that were in concordance with previous studies (Arun et al, 2013; Valdois et al, 2012).

The executive control interface with reading skill and language development is analyzed with association of nonsymbolic to symbolic performance (positive correlation, covariance of mazes, processing-speed and vocabulary) in HC which was modified in DDC group (no significant correlation of mazes, coding scores with vocabulary) and visual-search (picture-completion) performance was weakly associated to visuospatial reasoning, execution and reading performance (alphabet errors). Negative correlation of picture completion TQ scores and erroneous alphabets suggested that higher performance for visual-search was associated with lesser reading errors. These associations in two groups establish the interface of non-symbolic and symbolic performances.

Such Interface from the orthographic-visual analyzer (sublexical-lexical route) activates phonological lexicon (stored phonology), that in-turn activates phonological output buffer of speech production (Lukov et al, 2015) in efficient reading (Easson et al, 2020). In English monosyllabic minimal pairs, majority (84.2%) of the typical readers and only 44% DDC perceived 12-18 pairs correctly without any hearing loss. It is explained as poor categorical speech perception (Kronschnabel et al, 2014; Paul et al, 2006; Robertson et al, 2009; Valdois et al, 2008; Ziegler et al, 2009; Ziegler and Goswami, 2005) and/or poor auditory discrimination abilities (Law et al, 2014; Lorusso et al., 2014; Goswami et al, 2011) due to effortful, slow temporal sampling in auditory cortex (Ahmad, 2014; Goswami et al, 2011; Richards and Berninger, 2008) without any deficits of acoustic perception in pitch, rhythm and timbre (Grube et al, 2014).

In present study, non-significant differences in Hindi auditory discrimination (mean) and score distribution in two groups but significant difference (t = 1.861; p<0.05) and distribution for English monosyllabic minimal pairs (Figure 2,iv) emphasize the need to explore language orthographic depth (Nash et al, 2017; Suárez-Coalla et al, 2020). In both the groups (HC and DDC), minimum auditory discrimination scores were higher in Hindi (12-14 correct pairs in controls; 10-12 pairs dyslexics) as compared to English (10-12 correct pairs in controls; 4-6 pairs in dyslexics) (Figure 2,iv). Elucidated as learning ease of shallow orthography (closely matched morpho-semantic parameters; Nag and Snowling, 2013; Nash et al, 2017) and the daily use of language (sound mapping consistency) that is recognized in study participants as better auditory skill and vocabulary knowledge (McBride, 2016; Meschyan and Hernandez, 2006).

Thus the poor letter/ digit string processing and reading in dyslexia might not be only due to symbolic processing (phonological - Lobier et al, 2011) or letter-sound integration deficits (Nash et al, 2017) but comprise of perceptual, spatial attention allocation, and/or distractor-inhibition (Figure 2,v) at behavioural, cognitive and neural level (Al Dahhan et al, 2020; Alstad et al, 2015; Bellocchi and Leclercq, 2021; Bosse et al, 2007; Easson et al, 2020; Kronschnabel et al, 2014; Lobier et al, 2011; Pammer, 2014; Peyrin et al, 2011; Peyrin et al, 2012; Valdois et al, 2012; Xiaotong et al, 2012). Similarly, the poor execution into writing skill may implicate not only phonological-to-orthographic mapping but nosymbolic visual-visuospatial interface (Bellocchi and Leclercq, 2021; Bosse et al, 2007; Lobier et al, 2011; Nash et al, 2017; Pammer, 2014).

Present study has limitations of unequal sample size, indirect correlation of brain and behavioural measures. Although the study show higher performance variability in dyslexic group but the post-facto sub-grouping might not have yielded intended results for the inference (Bishop, 2013). Due to high variability in dyslexia it is recommended to analyze at individual or subgroups levels especially for therapeutic managements (Alstad et al, 2015; Roach and Hogben, 2007).

## 5. Conclusion

Reading is a complex integrative function of non-symbolic to symbolic processing and to unfold this mechanism four experiments were done in present study. Experiment-1 focused on pictorial visual-perception, reasoning and execution (picture completion, object assembly and block-design subtest); experiment-2 non-symbolic to symbolic processing (pattern recognition, coding, reading); experiment-3 symbolic execution (writing); and experiment-4 phonological awareness (auditory discrimination). In present study dyslexic readers have modified BOLD activation in parieto-frontal areas and higher activation in occipito-temporal regions (ventral stream) with performance variability in visual perception. Differences in visuospatial attention, distractor-inhibition and visuo-motor execution at presymbolic level in dyslexia with concomitant problems at symbolic level and at natural text reading (bigger string) suggests shared mechanism in non-symbolic and symbolic processing. Association of visual perception, visuospatial performance and executive control with vocabulary and reading errors further emphasize that these differences might not be limited to phonological and/or symbolic processing deficits only.

